# A *Rhodopseudomonas* strain with a substantially smaller genome retains the core metabolic versatility of its genus

**DOI:** 10.1101/2024.10.21.619483

**Authors:** Yasuhiro Oda, William C. Nelson, William G. Alexander, Stella Nguyen, Robert G. Egbert, Caroline S. Harwood

## Abstract

*Rhodopseudomonas* are a group of phototrophic microbes with a marked metabolic versatility and flexibility that underpins their potential use in production of value-added products, bioremediation and plant growth promotion. Members of this group have an average genome size of about 5.5 Mb, but two closely related strains have genome sizes of about 4.0 Mb. To identify the types of genes missing in a reduced genome strain, we compared strain DSM127 with other *Rhodopseudomonas* isolates at the genomic and phenotypic levels. We found that DSM127 can grow as well as other members of the *Rhodopseudomonas* genus and retains most of their metabolic versatility, but it has many fewer genes associated with high affinity transport of nutrients, iron uptake, nitrogen metabolism and biodegradation of aromatic compounds. This analysis indicates genes that can be deleted in genome reduction campaigns and suggests that DSM127 could be a favorable choice for biotechnology applications using *Rhodopseudomonas* or as a strain that can be engineered further to reside in a specialized natural environment.

**IMPORTANCE:** *Rhodopseudomonas* are a cohort of phototrophic bacteria with broad metabolic versatility. Members of this group are present in diverse soil and water environments and some strains are found associated with plants and have plant growth promoting activity. Motivated by the idea that it may be possible to design bacteria with reduced genomes that can survive well only in a specific environment or that may be more metabolically efficient, we compared *Rhodopseudomonas* strains with typical genome sizes of about 5.5 Mb to a strain with a reduced genome size of 4.0 Mb. From this we concluded that metabolic versatility is part of the identity of the *Rhodopseudomonas* group, but high-affinity transport genes and genes of apparent redundant function can be dispensed with.

## INTRODUCTION

*Rhodopseudomonas palustris* is a metabolically versatile alpha-proteobacterium that is a model for how bacteria integrate the functioning of diverse metabolic modules of aerobic respiration, carbon dioxide fixation, nitrogen fixation, and anaerobic photophosphorylation with organic compounds as carbon sources (1–5). It also produces hydrogen gas and is active in biodegradation (5–8). This metabolic versatility underpins increasing efforts to develop *R. palustris* for a variety of biotechnological applications including production of hydrogen gas and value-added products, as well as in bioremediation and plant growth promotion (9–12). It also explains the distribution of *Rhodopseudomonas* in diverse environments including swine sewage wastewater, rice paddy soils and high-altitude lagoons (13–15).

Most fundamental work on *R. palustris* has focused on two strains, CGA009 and TIE-1 (5, 16), but other *Rhodopseudomonas* with expanded metabolic versatility in anaerobic aromatic compound degradation, anaerobic growth in dark, light-harvesting and heavy metal detoxification have been described (15, 17, 18). Strains of *Rhodopseudomonas* that have been sequenced have genomes of between 4.9 and 6.0 Mb with an average genome size of 5.5 Mb. However, two strains with substantially smaller genomes of about 4.0 Mb have been isolated. One of these, strain DSM127, came from a pond, probably in Germany. The other strain, JSC-3b, was isolated from water in a canal adjacent to a vegetable field in Changsha, China (19).

Here we assembled a finished genome of the small genome strain DSM127 and compared its gene inventory and physiology to those of 38 other *Rhodopseudomonas* strains (Table S1). Our goal was to identify the types of functions that are missing from a free-living bacterium that has a relatively small genome compared to other members of its cohort. Such information may be useful in informing the design of genome reduction strategies to restrict the environmental persistence niche of bacteria or to engineer them for specialized metabolic functions.

## RESULTS

### Pangenome comparison of eleven Rhodopseudomonas stains

The set of nine strains analyzed by Lo et al. (14) is a useful comparator set for genome alignments, because it, along with strains DSM127 and JSC-3b, represents the breadth of phylogenetic diversity within the *Rhodospeudomonas* clade (15) (Fig. S1). A pangenome comparison highlights that there is a set of core genes shared by eleven strains, along with strain-specific genes (Fig. 1). We note that the genomes of strains JSC-3b and DSM127 are similar, with an average nucleotide identity of greater than 96.5%. For the remainder of this paper, we focus only on DSM127.

**FIG. 1.**
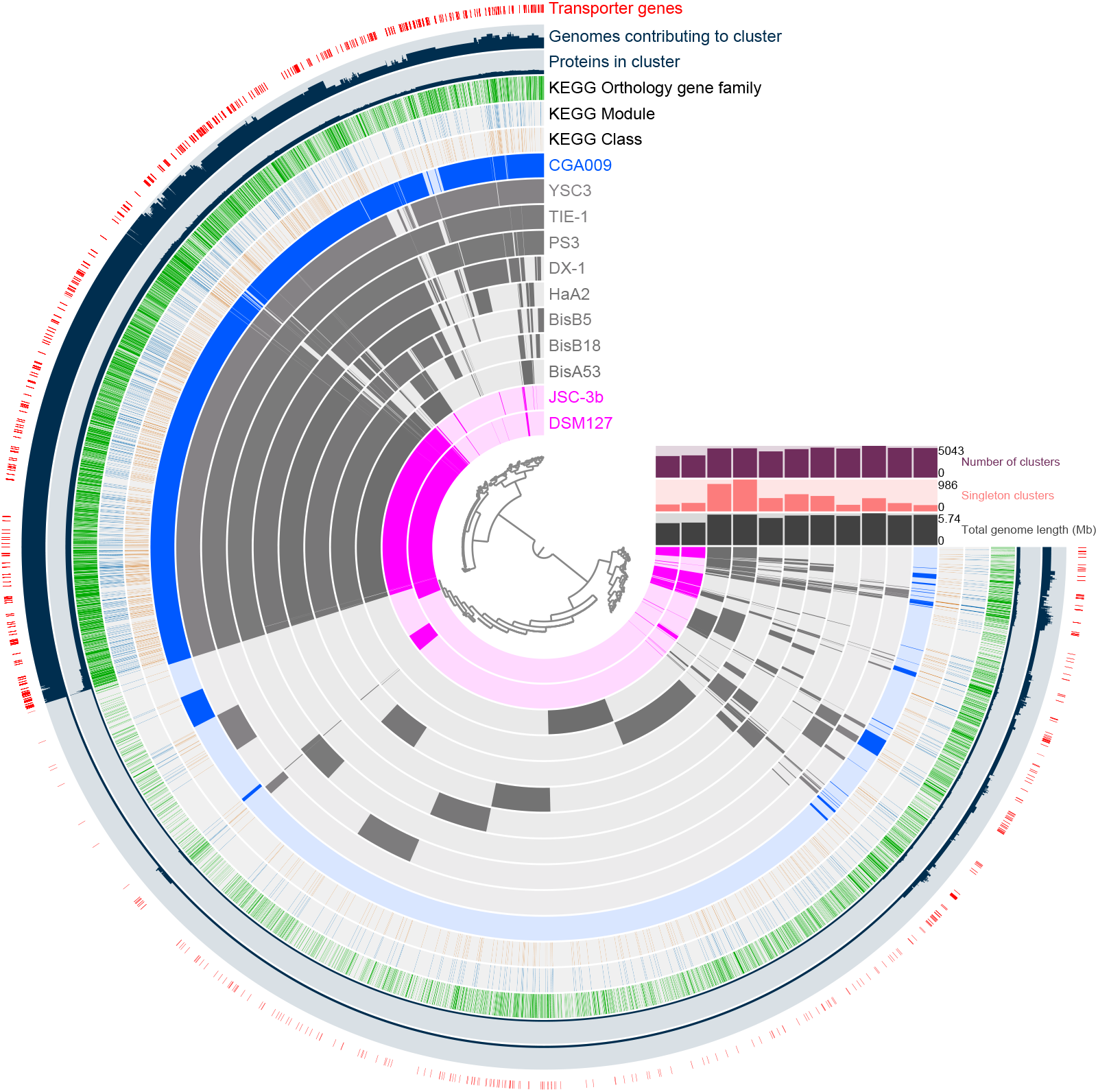
A pangenome comparison of 11 *Rhodopseudomonas* strains. The central dendrogram indicates hierarchical clustering of the proteins based on genome distribution profile. Layers from outer to inner: 1) predicted transporter genes (red); 2) number of genomes represented in cluster; 3) number of proteins assigned to cluster; 4) clusters with a KO family assignment (green); 5) clusters with a KEGG module assignment (blue); 6) clusters with a KEGG reaction class assignment (brown); 7 through 17) cluster membership by each of the Rhodopseudomonas strains as indicated. Bar plots: ‘Number of clusters’ – number of clusters to which proteins from the genome have been assigned; ‘Singleton clusters’ – count of protein clusters where all members are from this genome; ‘Total genome length (Mb)’ - nucleotide length of each genome.

### Comparison to strain CGA009

A comparison of protein-encoding genes of DSM127 with CGA009 shows that they have 2,536 orthologous genes (excluding paralogous genes), which is 52% of the genes in CGA009 and 71% of the genes in DSM127 (Tables S2a and S2b). There are 1,801 out of 4833 genes that are unique to CGA009 (Table S2c) and 840 out of 3,566 genes that are unique to DSM127 (Table S2d). About 16% of the genes that the strains have in common are annotated as hypothetical, conserved hypothetical, or genes of unknown function. About 39% of genes unique to CGA009 and 47% of the genes unique to DSM127 are annotated as hypothetical, conserved hypothetical, or genes of unknown function.

### Major physiological characteristics of strain DSM127

Microscopic examination of DSM127 showed that its cells have a dumbbell shaped morphology that is characteristic of CGA009 and other *Rhodopseudomonas*. DSM127 grew slightly faster than CGA009 in light under anaerobic conditions with acetate as the carbon source, but its growth rate in nitrogen-fixing conditions was slightly slower. DSM127 also had lower nitrogenase activity and produced less hydrogen than CGA009 (Table 1). When incubated anaerobically in light, strain CGA009 has a remarkable ability to survive and maintain full viability for weeks and months in a growth-arrested state (20). We found that this characteristic, which is useful when using cells as biocatalysts (21), was shared by DSM127 (Fig. 2A).

**TABLE 1.**
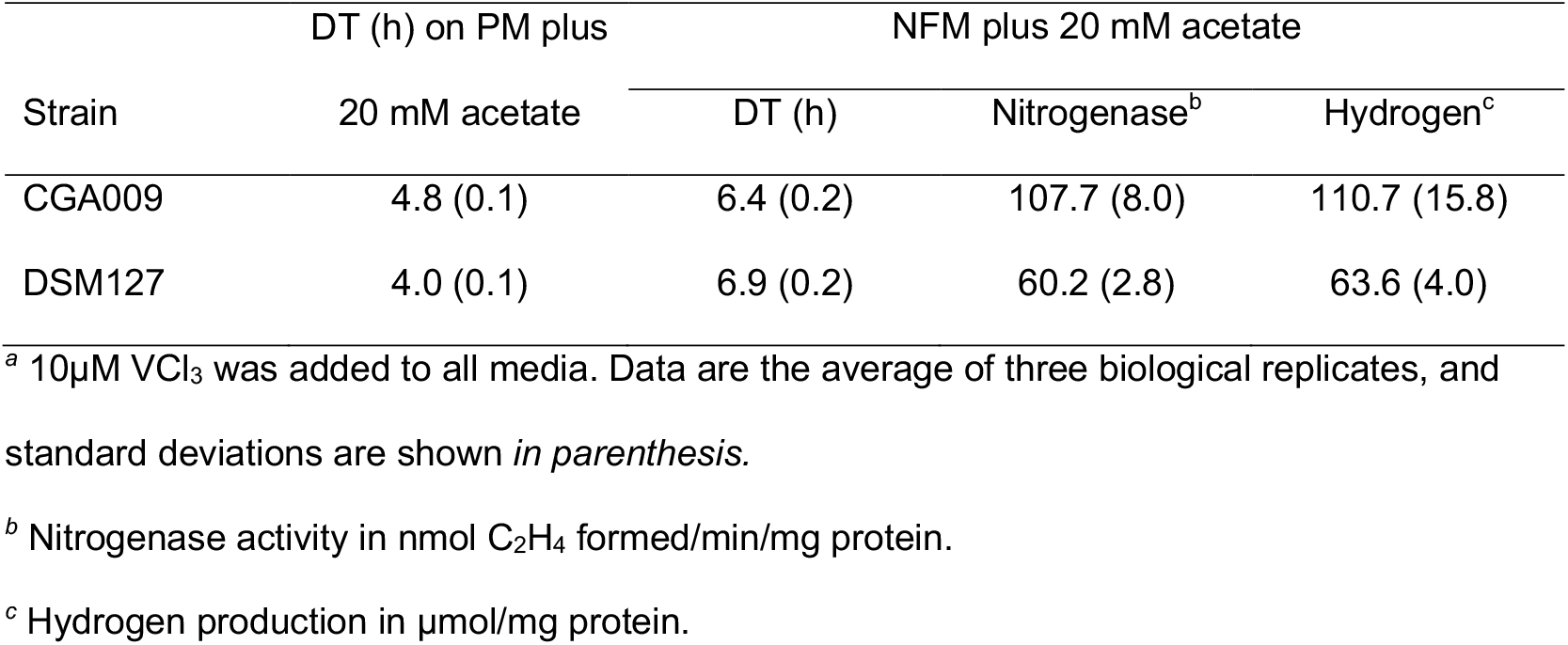
Doubling time (DT), nitrogenase activity, and hydrogen production from *Rhodopseudomonas* strains CGA009 and DSM127^*a*^.

**FIG. 2.**
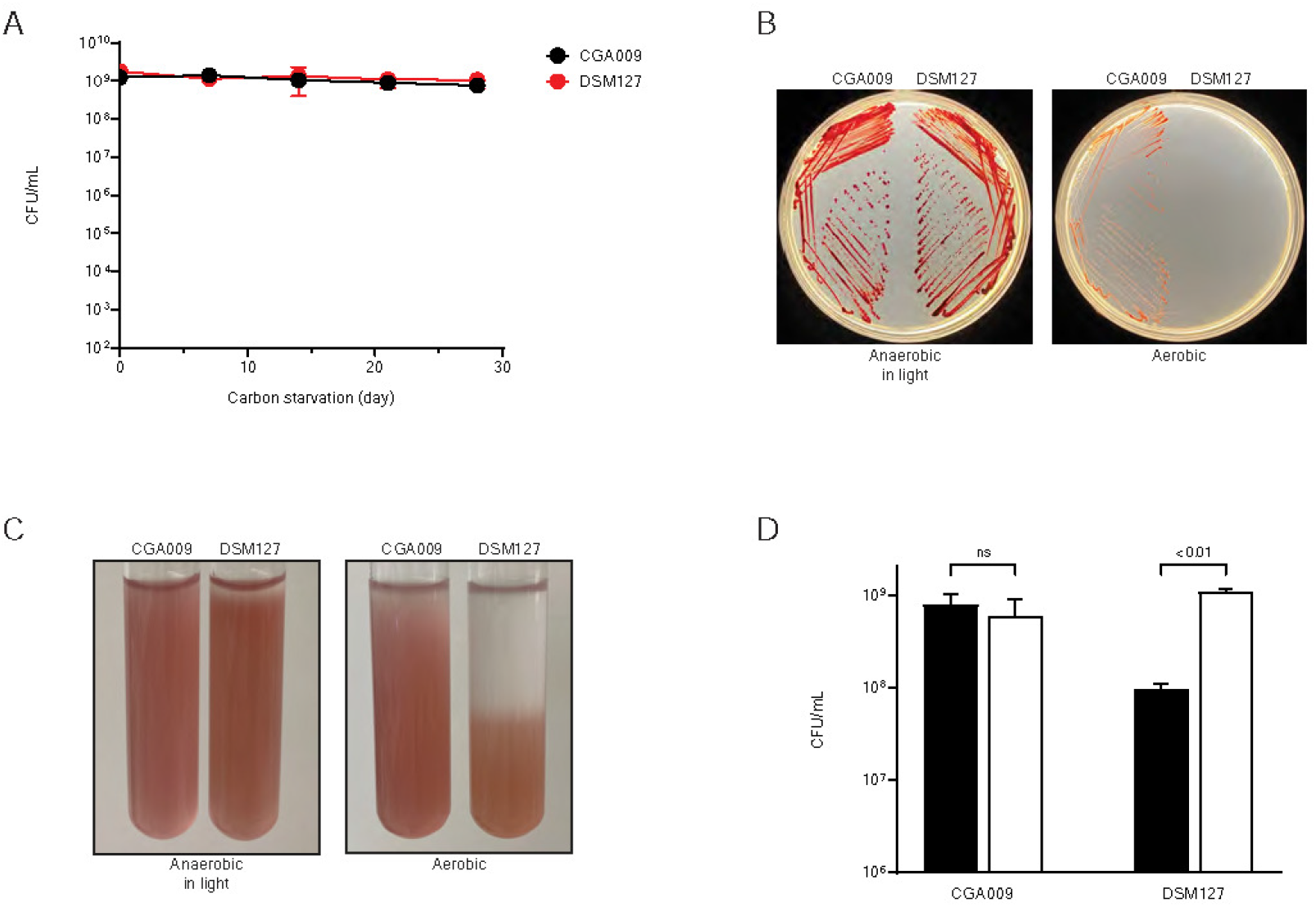
Comparison of physiological characteristics of strains CGA009 and DSM127. (A) Viability after growth arrest. Data are the average of three biological replicates, and standard deviations are shown as error bars. (B) Anaerobic and aerobic growth on CA agar (PM containing 20mM acetate supplemented with 0.2% Casamino acids and 0.5% yeast extract) for seven days. (C) Effect of oxygen on growth determined by agar shake assay (with 0.3% agar) after three days of cultivation. (D) Growth yield in low iron nitrogen-fixing medium (closed bars) and low iron nitrogen-fixing medium supplemented with 25µM FeSO4 (open bars). CFU were determined when culture reached the stationary phase of growth and the OD660 no longer increased. Data are the average of three biological replicates, and standard deviations are shown as error bars. Statistical significance was calculated using a t test: ns, not significant.

An obvious difference between DSM127 and other *Rhodopseudomonas* is that it does not form colonies on agar plates incubated in air (Fig. 2B). This was a surprise because from its genome sequence, DSM127 is predicted to grow aerobically. It shares with CGA009, *ccoNOQP* genes encoding ccb3 cytochrome c terminal oxidase and *coxBAEFCG* genes, encoding cytochrome aa3 oxidase. It also has genes for a cytochrome d ubiquinol oxidase that are present in some *Rhodopseudomonas* (17) (Table S3). We found that DSM127 cells grew in 0.3% agar a few cm below the air-agar interface in test tubes that were loosely capped and incubated in air (Fig. 2C). This indicates that DSM127 can grow microaerobically but is sensitive to atmospheric levels of oxygen. The basis of this apparent oxygen sensitivity is unclear, as DSM127 is not missing common oxidative stress tolerance genes, including catalase and superoxide dismutase genes and it has genes for an alkyl hydroperoxidase reductase (Table S3).

### Gene inventories

Core genes of *Rhodopseudomonas*, including strain DSM127, include those for nitrogen-fixation, photophosphorylation, oxidative phosphorylation, carbon source use, inorganic compound oxidation and carbon dioxide fixation (Table S4a). All *Rhodopseudomonas* have a conserved photosynthesis gene cluster that includes genes encoding reaction center proteins, light harvesting 1 polypeptides and carotenoid and bacteriochlorophyll biosynthesis enzymes (1). This cluster is intact in DSM127 except that one gene involved in regulation (*ppsR2)* (22) and several copies of bacteriophytochrome genes are missing. Many *Rhodopseudomonas* strains, including DSM127 have light harvesting 4 (LH4) genes that are regulated by a redox-sensing signal transduction system (18, 23). LH4 systems efficiently harvest light at low intensities and allow cells to grow in very low light. All *Rhodopseudomonas* including DSM127, encode both type I (*cbbSL*) and type II (*cbbM*) ribulose bisphosphate carboxylases for carbon dioxide fixation (24), as well as uptake hydrogenase genes and thiosulfate oxidation genes that are required for use of hydrogen and thiosulfate as electron donors for carbon dioxide fixation (25, 26).

Three global regulatory systems; FixLJ-K, RegRS and AadR, coordinate expression of genes for diverse metabolic modules, many of which make demands on the cellular electron supply in *Rhodopseudomonas*. In strains CGA009 and TIE-1, these systems respond to oxygen (FixLJ-K), intracellular redox status (RegRS) and anaerobiosis (AadR) (27–30). DSM127 encodes FixLJ-K, RegRS and AadR, but has only a subset of the genes known to be controlled by these regulators. It does not have, for example, genes for anaerobic benzoate and 4-hydroxybenzote degradation that are controlled by AadR, or the *pioABC* genes for anaerobic growth by iron oxidation that are regulated by FixK (27, 29). It also lacks many of the iron uptake genes regulated by RegRS (28).

### Strain DSM127 has less metabolic depth than other *Rhodopseudomonas* strains

When we compared carbon source use by DSM127 and CGA009 we found that both strains grew well on a variety of fatty acids including acetate, butyrate, caproate, glutarate, itaconate, citramalate, and citraconate. Both strains also grew on the amino acid L-leucine, but neither strain grew on other L-amino acids including alanine, glycine, valine, serine, methionine, tyrosine, arginine, tryptophan, asparagine or proline. A defining trait of *Rhodopseudomonas* is the use of dicarboxylic acids for growth (31). We found that, like CGA009, DSM127 grew well on dicarboxylic acids of 7 carbons (pimelate), 9 carbons (azelate), 10 carbons (sebacate) and 12 carbons (dodecanedioate). DSM127 has most of a cluster of *pim* genes in CGA009 that are required for optimal growth on pimelate and azelate (31).

DSM127 lacks genes for anaerobic aromatic compound degradation that have been described in CGA009 (32–34), and this was one category of potential substrates that it was unable to use for anaerobic growth. DSM127 did not grow on benzoate, 4-hydroxybenzoate, p-coumerate, cinnamate or caffeate; compounds that all support the growth of CGA009. Anaerobic growth on aromatic compounds has been considered a hallmark of the *Rhodopseudomonas*, but one previously described strain, HaA2, also lacks genes for this trait (17).

A COG analysis of the 38 *Rhodopseudomonas* strain set (excluding the small genome strain JSC-3b) showed that DSM127 has fewer genes in all COG categories (Fig. S2), indicating a very broad loss of functionality. Examining individual COG subcategories, we observed that DSM127 has about half the number of genes encoding enzymes for acyl-CoA synthetases (COG0318) and enoyl-CoA hydratases (COG1024) than other *Rhodopseudomonas*. These categories of proteins are important for catabolism of organic compounds that have carboxyl groups, including fatty acids, aromatic acids, and dicarboxylic acids (31–33) (Table S4a).

*Rhodopseudomonas* strains encode relatively large numbers of genes annotated as ABC transporters and TRAP transporters for the uptake of various nutrients including iron, C-4 dicarboxylates, oligopeptides, and aromatic compounds. These two types of transporters have associated genes for high affinity solute binding proteins, which are also known as periplasmic binding proteins (35–37). Strain CGA009 and other *Rhodopseudomonas* strains encode about 120 solute binding proteins (COG description containing “periplasmic component” in Table S4a), suggesting that they can scavenge low amounts of diverse nutrients from their environment.

DSM127 encodes only about 50 solute binding proteins, and we hypothesize that this translates into a reduced ability to compete for scarce nutrients relative to other *Rhodopseudomonas*.

DSM127 has a narrower inventory of nitrogen utilization genes than other *Rhodopseudomonas*. A COG category comparison of DSM127 with the nine other best characterized *Rhodopseudomonas* strains (14) (Table S4b) shows that all strains contain three nitrogen regulatory PII proteins (COG0347: called *glnB, glnK1* and *glnK2*) whereas DSM127 has *glnB* and one *glnK* gene. DSM127 is the only one of these strains that does not encode urease for urea utilization. All *Rhodopseudomonas* can convert nitrogen gas to ammonia by nitrogen-fixation catalyzed by nitrogenase enzymes. Three forms of nitrogenase isozymes; molybdenum nitrogenase, vanadium nitrogenase and iron-only nitrogenase have been characterized, but molybdenum nitrogenase is the most widespread and efficient of these and is found in all nitrogen fixing bacteria (38). Most strains of *Rhodopseudomonas* encode iron-only nitrogenase in addition to the molybdenum enzyme and some strains, including strain CGA009, also have vanadium nitrogenase. DSM127 joins strains BisB5 and HaA2 in encoding only molybdenum nitrogenase (17).

*Rhodopseudomonas* requires large amounts of iron for synthesis of its electron carriers and nitrogenase enzyme. We have found that it is difficult to starve strain CGA009 for iron in the laboratory and this is reflected in its genome, which encodes 24 outer membrane receptor proteins (COG1629), most of which are predicted to be involved in transport of ferric iron [Fe(III)]. By contrast DSM127 has just six outer membrane receptor proteins. Strains CGA009, TIE-1, DX-1 and HaA2, but not strain DSM127, have genes for rhodopetrobactin siderophores that have high affinity for Fe(III) (39). Ferrous iron [Fe(II)] is more soluble than ferric iron and can be directly transported by single subunit transporters. Strains CGA009, TIE-1, and PS3 have genes for six Fe(II) transporters and DSM127 encodes a subset of four of these (40).

Strain CGA009 synthesizes ferrosomes from six *fez* genes (RPA2333-2338) (41). DSM127, however, lacks a complete gene set for these iron-containing particles, which are proposed to serve as an iron-storage organelle in anaerobiosis. As might be expected from its reduced number of iron transport and acquisition genes, we found that that DSM127 grew to a lower yield in low iron medium compared to strain CGA009 (Fig. 2D).

### The DSM127 genome is enriched in phage defense systems

DSM127 has two clusters of CRISPR-associated protein genes. Most strains, including strains CGA009 and TIE-1, lack CRISPR-CAS systems, but a few strains have one system. DSM127 also has at least two toxin-antitoxin gene sets and encodes a type I restriction-modification system (Table S3) which, like the CRISPR-Cas systems, are probably involved in phage defense.

## DISCUSSION

Our phenotypic and genomic analyses shows that the reduced genome strain DSM127 retains the distinctive combination of core characteristics that sets the *Rhodopseudomonas* clade apart from other phototrophic bacteria and from other members of the family *Bradyrhizobiaceae*.

Phenotypic characteristics that are specific to DSM127 relative to other *Rhodopseudomonas* are its inability to grow in fully aerobic conditions, its relatively poor ability to grow under conditions of iron depletion, and its inability to use aromatic compounds as carbon sources. From its genome sequence, we can see that DSM127 lacks the metabolic depth characteristic of *Rhodopseudomonas*. It encodes just one of three nitrogenase isozymes, it lacks genes for use of reduced iron as an electron donor for photoautotrophic growth and has many fewer outer membrane receptor transport systems and solute binding proteins than other *Rhodopseudomonas* strains. These features suggest that DSM127 occupies a narrower range of natural microenvironments than its relatives with larger genomes. Further supporting this hypothesis is the lack of strain-specific genes of known function in DSM127, whereas other *Rhodopseudomonas* strains have strain-specific metabolic versatility that is conferred by auxiliary genes (14, 15, 17). These auxiliary genes are predicted to enable some strains to thrive in specific microenvironments, including environments with high concentrations of heavy metals or deeper in soils where little to no light penetrates.

This comparative study suggests that if one wanted to engineer a *Rhodopseudomonas* strain or another kind of bacterium to have a smaller genome, then appropriate targets for deletion might be genes for transporters, especially high affinity transporters and other genes such as vanadium- and iron-nitrogenase genes that are used only under specialized conditions (38, 42). One can hypothesize that removal of transporter genes and genes of apparent redundant functions from the genome of a bacterial strain would thwart that strain’s ability to thrive in some environments relative to its wild-type parent. This could be a first step towards engineering a bacterium that is restricted to a specific environmental niche. An example of a specific niche might be plant root rhizospheres. Some *Rhodopseudomonas* strains have plant-growth promoting activity (11, 43, 44) and it might be useful to engineer such a strain to grow only when in close physical association with a specific host plant.

Our analysis raises the question of how strain DSM127 evolved to have a small genome.

While it is out of the scope of this study to try to address this question, a comparison of the DSM127 and CGA009 genomes shows that the two genomes line up quite well in terms of the orders of their homologous gene sets (Table S2). However, DSM127 is missing blocks of contiguous genes that are present in CGA009. Similarly, DSM127 has acquired blocks of contiguous genes. The DSM127 genome contains few pseudogenes, and its genome does not appear to be degraded. This suggests a mechanism of episodic gain or loss of genomic islands (45), allowing for rapid changes in genome size and metabolic capability.

## MATERIALS AND METHODS

### Bacterial strains and growth conditions

*Rhodopseudomonas* strains were routinely cultivated in PM medium (46) supplemented with Wolfe’s vitamin solution (5 ml per liter of medium) (47) and incubated at 30°C under anaerobic conditions in light. Unless otherwise mentioned, 20 mM acetate was used as the sole carbon source. Cells were grown under nitrogen-fixing conditions in PM without ammonium sulfate (NFM) with a nitrogen gas, and nitrogenase activity and hydrogen production were measured as previously described (42). NFM with no added FeSO4 or 25 µM FeSO4 was used for low and high iron conditions (48). Cell survival assays during carbon starvation were performed as previously described (49).

### Whole genome sequencing

The genome of strain DSM127 was sequenced with PacBio RS II (at CD Genomics, Shirley, NY, USA) and Illumina HiSeq 2000 platforms. 2.1 Gbp of reads with an N50 of 9.9 kbp were obtained from the PacBio run. This read set was filtered by removing all reads under 1 kbp, then Trycycler v0.5.3 (https://github.com/rrwick/Trycycler) was used in conjunction with the Raven v1.5.3 (50), Miniasm/Minpolish v0.3-r179 (51), and Flye v2.9 assemblers (52) to produce an assembly consisting of a single 3.8 Mbp chromosome made from a read depth of ~500x. This sequence was polished using Illumina HiSeq 2000 short reads downloaded from GenBank (SRX867298 and SRX867299) by first filtering each dataset via the USEARCH (53) fastq_filter feature with settings ‘-fastq_truncqual 19 -fastq_minlen 20’, producing a read pool at a coverage depth of ~130x. This pool was then used with Polypolish v0.6.0 (54) and BWA-MEM 0.7.17-r1188 (55), which changed seven bases from the initial Trycycler consensus sequence; all but ~900 bases were covered at 5x or greater by the short-read set. The final genome sequence was deposited to IMG database (https://img.jgi.doe.gov/) under IMG submission ID 274530.

### Comparative genome analysis

Reference genome sequences were downloaded from JGI IMG database (Table S1). Genomes were converted to Anvi’o (v7) databases using gene predictions from IMG (56) and KEGG Ortholog family annotation was calculated using anvi-run-kegg-kofams. Genome comparison was performed using anvi-pan-genome and visualized with anvi-display-pan. Presence/absence analysis (Table S2a) between CGA009 and DSM127 was based on OrthoFinder (57) results and TIGRfam (58) membership integrated using boutique code. COG annotations were performed as part of the IMG annotation pipeline (59).

### Carbon source utilization assay

A technique called auxanography was used to test for growth of strains DSM127 and CGA009 on various carbon sources under anaerobic conditions in light (60). Cells were grown in PM with 10 mM acetate supplemented with Wolfe’s vitamin, harvested, washed and diluted to an OD660 of 0.5 in PM without carbon source. Cell suspension (50 ml) was added to 1 liter PM medium containing 0.5% (wt/vol) melted Gelzan™ CM Gellan Gum (CP Kelco) and the mixture was immediately poured into petri dishes. Each carbon source to be tested was added either as a small amount of liquid or a small amount of solid as a spot on the solidified surface of the gelled medium near the periphery of the petri dish. Petri dishes were placed in GasPak EZ Anaerobe Gas Generating Pouches (BD) and incubated at 30°C under anaerobic conditions in light for seven days. An arc of growth was observed around the point of carbon compound addition on the petri dish for those substrates that supported growth.

### Agar-shake assay to test growth in an oxygen gradient

Cells grown in PM with 20 mM acetate were diluted to an OD660 of 1.0 in PM without carbon source, and 1 ml of this cell suspension was added to sterile 50 ml tubes containing 20 ml PM, 20 mM acetate and 0.3% Difco Agar Noble (BD) that were held in a 60°C water bath. After mixing, eight ml of this agar mixture was added to sterile 16 X100 mm test tubes and Hungate tubes. For aerobic conditions, a test tube cap was loosely placed on test tubes. For anaerobic conditions, Hungate tubes were flushed with nitrogen gas for 30 min before being sealed with butyl stoppers. Aerobic and anaerobic tubes were incubated in front of two 60 W light bulbs.

## Data Availability

The complete genome sequence of strain DSM127 was deposited to IMG database (https://img.jgi.doe.gov/) under IMG submission ID 274530.

## ACKNOWLEDGEMENTS

This research was supported by the Secure Biosystems Design Program, US Department of Energy, Office of Science, Biological and Environmental Research, as part of the PNNL Persistence Control (PerCon) Science Focus Area. Pacific Northwest National Laboratory is operated by Battle for the US Department of Energy under Contract DE-AC05-76RL01830.

## SUPPLEMENTAL MATERIAL

**FIG. S1.**
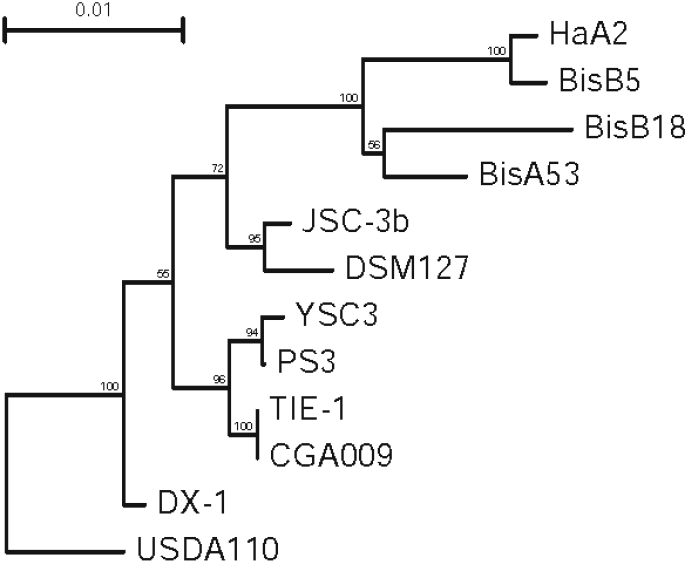
Phylogenetic relationships of *Rhodopseudomonas* strains based on 16S rRNA gene sequences. Bootstrap values (100 replicates) are given at branch points (only showing values of > 50). Bar represents substitutions per site. *Bradyrhizobium diazoefficiens* USDA110 was used to root the tree.

**FIG. S2.**
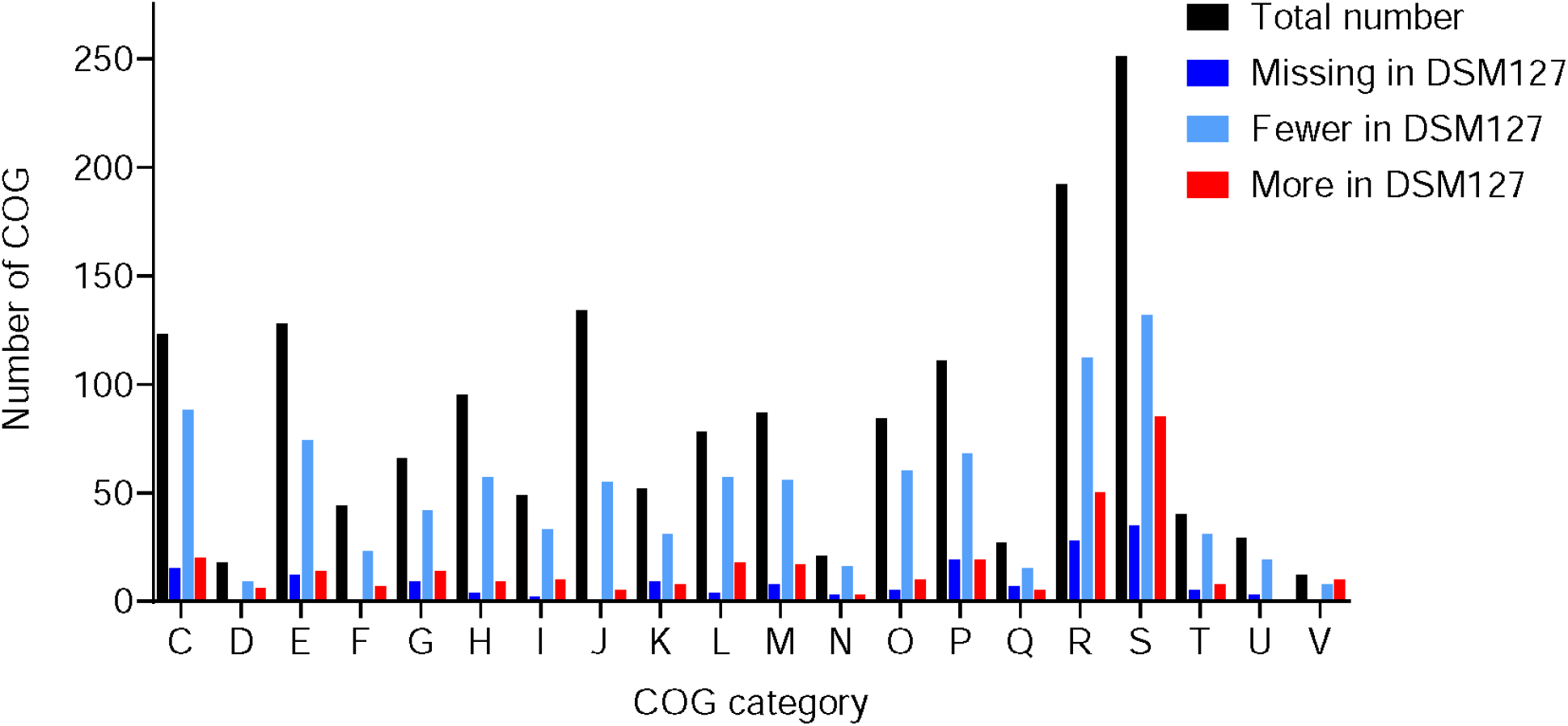
COG category analysis of DSM127 against 37 other *Rhodopseudomonas* strains. COG categories: C, Energy production and conversion; D, Cell cycle control, cell division, chromosome partitioning; E, Amino acid transport and metabolism; F, Nucleotide transport and metabolism; G, Carbohydrate transport and metabolism; H, Coenzyme transport and metabolism; I, Lipid transport and metabolism; J, Translation, ribosomal structure and biogenesis; K, Transcription; L, Replication, recombination and repair; M, Cell wall/membrane/envelope biogenesis; N, Cell motility; O, Posttranslational modification, protein turnover, chaperones; P, Inorganic ion transport and metabolism; Q, Secondary metabolites biosynthesis, transport and catabolism; R, General function prediction only; S, Function unknown; T, Signal transduction mechanisms; U, Intracellular trafficking, secretion, and vesicular transport; V, Defense mechanisms. For this figure only, we first calculated the average number of gene counts for each COG using our 37 *Rhodopseudomonas* strain set (excluding DSM127) (Table S1) and compared with the gene count from DSM127. Those COGs that had an average count of less than one were excluded. See Table S4 for detailed COG analysis of each strain.

**TABLE S1**. Reference genome sequences used in this study (supplied as an Excel file).

**TABLE S2**. Gene inventory comparisons of *Rhodopseudomonas* strains CGA009 and DSM127 (supplied as an Excel file).

**TABLE S3**. Gene products and locus tags of some of the DSM127 genes described in this study (supplied as an Excel file).

**TABLE S4**. A COG analysis of *Rhodopseudomonas* strains (supplied as an Excel file).

